# Proteins as Statistical Languages: Information-Theoretic Signatures of Proteomes Across the Tree of Life

**DOI:** 10.64898/2025.12.31.697235

**Authors:** Enso O. Torres Alegre

## Abstract

Protein sequences are commonly interpreted through biochemical and evolutionary lenses, emphasizing structure–function relationships and selection in sequence space. Here we develop a complementary viewpoint: proteins as *statistical languages*—strings over a finite alphabet generated by constrained stochastic processes. We formalize intrinsic informational descriptors of protein ensembles, including composition entropy *H*_1_, adjacent mutual information *I*_1_, and separation-dependent information profiles *I*_*d*_. A null-model ladder (uniform, composition-matched i.i.d., and Markov-1) separates compositional effects from genuine positional dependence. We then evaluate these descriptors empirically across 20 UniProt reference proteomes spanning major clades, using protein-level bootstrap resampling and matched synthetic controls. Real proteomes consistently depart from composition-matched i.i.d. baselines and exhibit information profiles that remain elevated beyond the decay expected under first-order Markov surrogates, indicating dependencies beyond local transition statistics. Finally, a compressibility proxy (gzip) provides an orthogonal signature of redundancy relative to i.i.d. controls at matched composition. Together, these results support the view of proteomes as constrained statistical languages and provide model-agnostic fingerprints for comparing sequence ensembles.These signatures provide a lightweight diagnostic layer for comparing proteomes prior to mechanistic modeling

## 1 Introduction

Proteins are linear polymers over an alphabet of 20 amino acids that fold into three-dimensional structures and realize biological functions. Standard approaches interpret sequences through physicochemical constraints (e.g. stability and foldability) and evolutionary mechanisms operating in sequence space [2, 9, 12, 14]. Alongside these mechanistic views, protein sequences also display robust statistical regularities: non-uniform residue frequencies, context-dependent patterns, and correlations at multiple length scales [3, 5, 6, 16]. These properties suggest that proteins—viewed purely as strings—occupy a small and structured subset of the space of all possible strings.

Information theory and statistical linguistics quantify structure in symbolic sequences using entropy, mutual information, and compressibility [1, 8, 15]. Importantly, these tools detect constraints without requiring semantic interpretation. This motivates the central hypothesis explored here:

*Protein sequences can be treated as realizations of a statistical language: a constrained generative process over a finite alphabet*.

This hypothesis does *not* claim proteins are encrypted messages. Rather, it frames compositional bias and context dependence as information-theoretic signatures of constrained generation, enabling principled comparison between ensembles.

### Contributions

This paper (i) formalizes proteins as statistical objects and defines intrinsic informational measures; (ii) introduces a hierarchy of null models that separate composition from higher-order structure; (iii) defines information profiles across sequence separations; and (iv) proves a general entropy-deficit bound showing that rarity/viability constraints necessarily reduce achievable entropy. To demonstrate that these descriptors are not merely formal, we perform a cross-proteome analysis on 20 UniProt reference proteomes [17], spanning 10 biological categories (two proteomes each), using protein-level bootstrap resampling and two matched null controls (composition-matched i.i.d. and Markov-1). This empirical component tests whether real proteomes occupy distinct regions in (*H*_1_, *I*_1_) space and whether their information profiles *I*_*d*_ systematically exceed those of finite-memory surrogates.

### Scope

Our goal is deliberately model-agnostic: to quantify statistical structure in proteome sequence ensembles without requiring structural annotation, functional labels, or family alignments. The resulting signatures should be viewed as descriptive fingerprints rather than mechanistic explanations.

## 2 Protein Sequences as Statistical Objects

Let 𝒜 denote the amino acid alphabet with |𝒜| = 20. A sequence of length *L* is

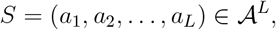

and the space of all finite-length sequences is 𝒮= ∪_*L≥*1_ 𝒜^*L*^. We treat an ensemble of sequences as samples from a probability distribution ℙ over 𝒮. For fixed *L*, write ℙ_*L*_ for the induced distribution on 𝒜^*L*^.

### Definition 2.1

(Composition-matched i.i.d. baseline). *Given a one-site marginal p*_1_ *over* 𝒜, *the composition-matched i*.*i*.*d. model on* 𝒜^*L*^ *is*

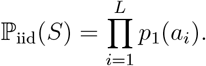

This baseline isolates structure beyond composition: any dependency between positions reduces entropy relative to the maximum-entropy distribution with the same one-site marginals [1, 11].

## 3 Amino Acid Usage and Entropy

### 3.1 Composition

Given an ensemble, define the marginal frequency

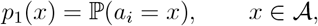

estimated by pooled counts across sequences. The compositional entropy is

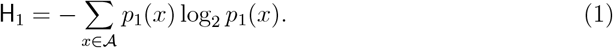

To compare composition against a reference *q* (e.g. uniform or a background), use Kullback– Leibler divergence

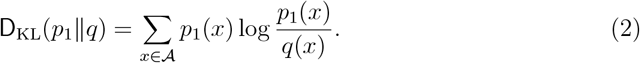

### 3.2 Sequence entropy and an extremal principle

Let H(*S*) denote the Shannon entropy of ℙ_*L*_ on 𝒜^*L*^:

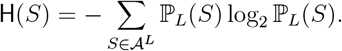

#### Proposition 3.1

(Maximum entropy at fixed one-site marginals)

*Among all distributions on* 𝒜^*L*^ *with fixed one-site marginals p*_1_, *the composition-matched i*.*i*.*d. model* 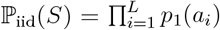 *maximizes* H(*S*).

*Proof*. Fixing one-site marginals defines a convex set of admissible distributions. The product measure is the unique maximizer because any statistical dependency introduces additional constraints that reduce entropy [1, 11].

Proposition 3.1 separates (i) compositional constraints from (ii) higher-order constraints encoded in reduced conditional entropies and nonzero mutual information.

## 4 Correlations and Information Profiles

### 4.1 Local (second-order) structure

Define adjacent joint frequencies

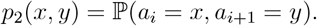

The conditional entropy is

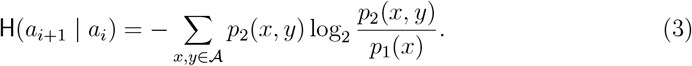

The adjacent mutual information is

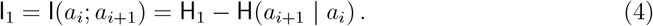

If residues are i.i.d. (given *p*_1_), then I_1_ = 0.

### 4.2 Length-scale dependence

For separation *d* ≥ 1, define

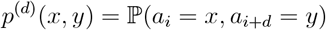

and mutual information

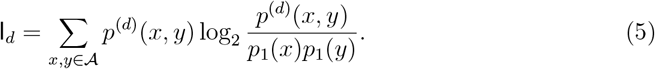

#### Definition 4.1

(Information profile)

*The mapping d* ↦ I_*d*_ *is the* information profile *of an ensemble. Its decay characterizes the range of dependencies (short-range vs. long-range)*.

#### Remark 4.1

*For a finite-order Markov source, dependencies are predominantly local; empirically, one often observes rapid decay of* I_*d*_ *with d once the relevant memory scale is exceeded. Slow decay or plateaus can arise from global constraints, repeats, or mixture structure, and do not by themselves imply semantic “meaning*.*”*

#### Lemma 4.1

(Independence and vanishing information profile)

*If* I_*d*_ = 0 *for all d* ≥ 1 *and the process is stationary with well-defined one-site marginal p*_1_, *then all pairs factorize:*

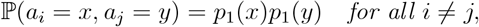

*i*.*e. the process is pairwise independent. In particular, any departure from i*.*i*.*d. behavior must yield* I_*d*_ *>* 0 *for at least one separation d*.

*Proof*. By definition, I_*d*_ = 0 implies *p*^(*d*)^(*x, y*) = *p*_1_(*x*)*p*_1_(*y*) for all *x, y* ∈ 𝒜 at that separation. If this holds for all *d* ≥ 1, then every pair of positions factorizes into the product of marginals, yielding pairwise independence.

## 5 Proteins as Statistical Languages

We formalize “language” in a statistical (not semantic) sense: a constrained set of admissible strings together with a generative distribution.

### Definition 5.1

(Statistical protein language)

*A* statistical protein language *is a pair* (ℒ, ℙ) *where* ℒ ⊆ 𝒮 *is a set of admissible sequences (constraints) and* ℙ *is a probability measure supported on* ℒ.

### 5.1 Null-model ladder

A principled comparison ladder is:

1. **Uniform random (UR)**: ℙ_UR_(*S*) = 20^−*L*^ on 𝒜^*L*^.
2. **Composition-matched i.i.d. (IID)**: ℙ_iid_(*S*) = Π_*i*_ *p*_1_(*a*_*i*_).
3. **Finite-order Markov (Mk)**: matches *k*-mer statistics (here *k* = 1).
4. **Maximum-entropy (ME)**: matches selected constraints while maximizing entropy [7].

Each rung provides a benchmark; deviations quantify additional structure.

### 5.2 Maximum-entropy grammars

Following Jaynes [7], maximum-entropy models provide a canonical “grammar” induced by constraints. Given constraint functions *f*_*m*_(*S*) with target averages, the ME distribution takes the exponential-family form

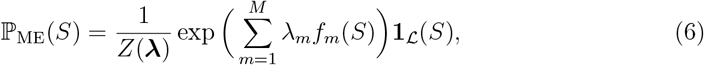

where *Z*(***λ***) normalizes.

#### Proposition 5.1

(Constraints induce dependencies)

*If* (6) *includes only one-site constraints, then* ℙ_ME_ *is composition-matched i*.*i*.*d. and* I_*d*_ = 0 *for all d. Introducing multi-site constraints generically yields* I_*d*_ *>* 0 *at corresponding separations*.

#### Remark 5.1

*Pairwise maximum-entropy models are closely related to Potts/DCA formulations used to capture covariation in protein families [3, 6, 18]*.

## 6 Empirical Data and Methods

### 6.1 Proteomes and preprocessing

We analyzed 20 proteomes downloaded from UniProt via the UniProt REST API, using UniProt reference proteome identifiers spanning 10 broad biological categories (two proteomes per category): mammals, birds, fish, amphibians, insects, plants, bacteria, archaea, filamentous fungi, and yeasts [17]. Protein sequences were parsed from FASTA format, filtered to retain only canonical amino acids (20-letter alphabet), and truncated by removing sequences shorter than a minimum length (here *L*_min_ = 80) to reduce instability from extremely short proteins. For computational feasibility, large proteomes were subsampled using a fixed maximum number of sequences per proteome; all analyses below should thus be interpreted as estimates on a representative proteome subsample.

### 6.2 Bootstrap protocol

For each proteome, we performed protein-level bootstrap resampling. Each bootstrap replicate consists of sampling proteins with replacement from the filtered proteome (sample size *n* = 200 proteins per replicate), preserving the empirical length distribution within that replicate. We used 150 bootstrap replicates per proteome. All reported summary statistics are means and standard deviations across bootstrap replicates.

### 6.3 Information-theoretic estimators

One-site marginals *p*_1_ were estimated by pooled residue counts. Adjacent joint frequencies *p*_2_ and separated joint frequencies *p*^(*d*)^ were estimated by direct counting. Mutual informations I_1_ and I_*d*_ were computed using plug-in estimators (base-2 logarithms; units are bits). The information profile was evaluated for separations *d* = 1, …, 25.

### 6.4 Matched null controls

For each bootstrap replicate, two controls were generated:

1. **IID control:** sequences generated by sampling residues i.i.d. according to the replicate-specific empirical *p*_1_, using the same length distribution as the real replicate.
2. **Markov-1 control:** sequences generated by a first-order Markov chain with initial distribution *p*_1_ and transition matrix estimated from the real replicate dipeptide counts *p*_2_, again using the same length distribution.

These controls isolate compositional effects from positional dependence and test whether observed dependencies extend beyond first-order transition statistics.

### 6.5 Compressibility proxy

As an orthogonal signature of redundancy, we computed an overhead-corrected gzip code length per character on a long concatenation of sampled sequences (with a fixed total amino-acid budget). This measure is a practical proxy for algorithmic redundancy rather than a formal entropy-rate estimator [4, 10].

### 6.6 Code and reproducibility

All analyses and figure generation are fully reproducible from a public Google Colab/Jupyter notebook available at GitHub repository (InTheoryGeneral.ipynb).

The notebook implements UniProt proteome retrieval via the REST API, preprocessing and filtering (*L*_min_ = 80), bootstrap resampling, IID and Markov-1 surrogate generation, entropy and mutual-information estimation, and gzip-based compressibility comparisons, reproducing Figures 1–4 from the same pipeline.

**Figure 1:**
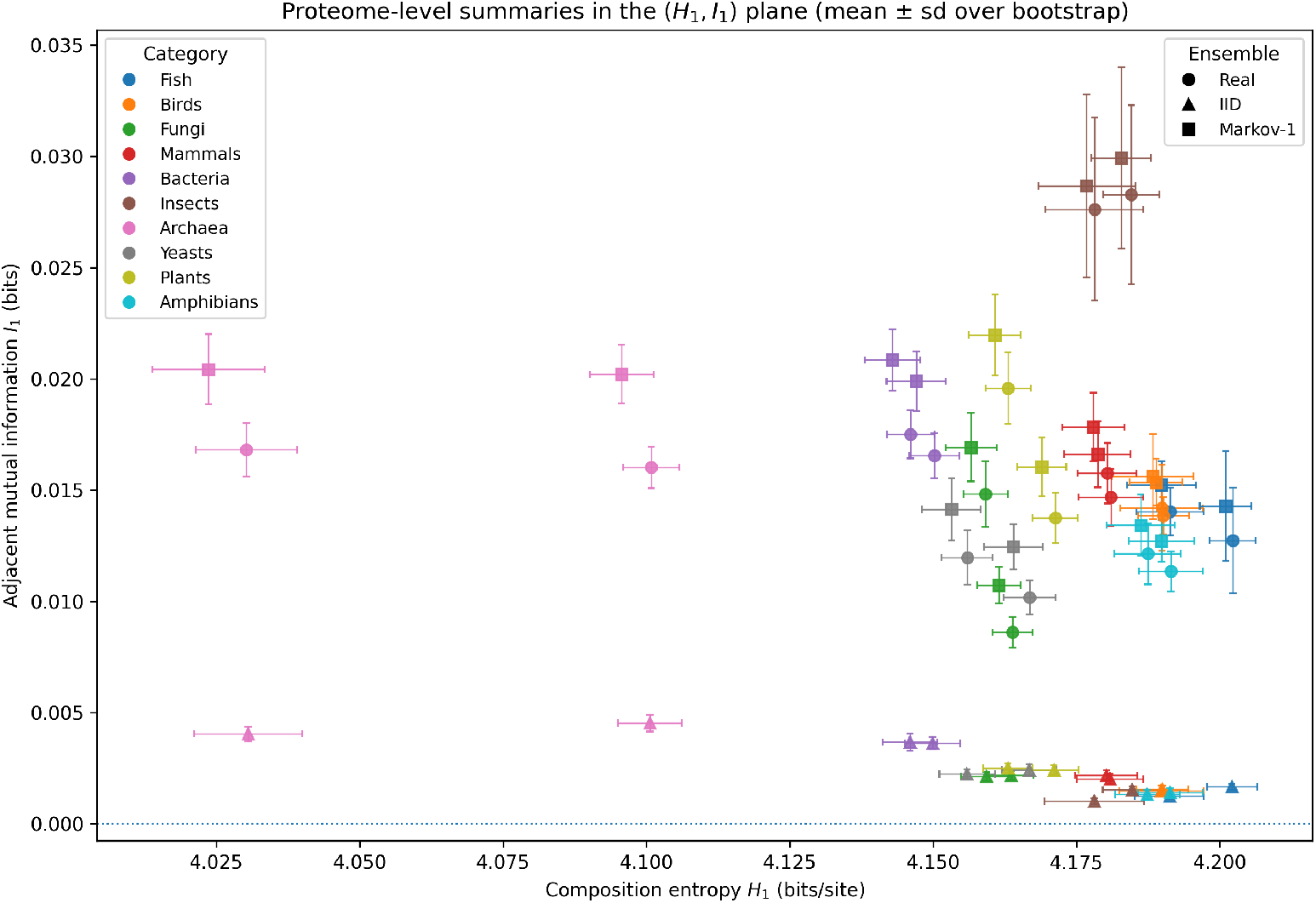
Proteome-level summaries in the (*H*_1_, *I*_1_) plane (mean ± s.d. across bootstrap replicates). Colors indicate broad biological categories and marker shapes indicate the ensemble (Real, IID, Markov-1). IID controls preserve one-site composition *p*_1_ while destroying positional dependence, yielding *I*_1_ ≈ 0. Markov-1 controls preserve adjacent dipeptide statistics and reproduce local dependence by construction. Real proteomes are systematically displaced from IID baselines, indicating nontrivial ordering constraints beyond composition alone.

## 7 Entropy Deficit Under Viability Constraints

We state a general result requiring no explicit folding or functional model.

### Definition 7.1

(Viable set and rarity fraction). *Let* 𝒱_*L*_ ⊆ 𝒜^*L*^ *be the set of “viable” sequences of length L under some broad constraint. Define the rarity fraction*

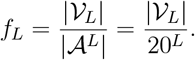

### Theorem 7.1

(Entropy deficit bound). *For any distribution* ℙ *supported on* 𝒱_*L*_,

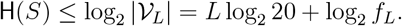

*Equivalently*,

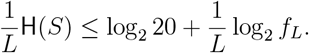

*Proof*. Entropy is maximized by the uniform distribution on the support. The uniform distribution on 𝒱_*L*_ has entropy log_2_ |𝒱_*L*_|, which upper-bounds any supported distribution [1].

### Corollary 7.1.1

(Rarity implies unavoidable structure)

*If f*_*L*_ *decays exponentially with L (e*.*g. f*_*L*_ ∼ 2^−*αL*^ *with α >* 0*), then no viable-sequence ensemble can be simultaneously close to uniform composition* and *close to i*.*i*.*d. ordering in the sense that both* H_1_ ≈ log_2_ 20 *and* I_*d*_ ≈ 0 *for all d hold asymptotically*.

### Remark 7.1.

*Corollary 7.1.1 provides a clean conceptual link: if viability is rare, statistical structure is unavoidable and must appear either as compositional bias, correlations, or both*.

## 8 Results: Cross-Proteome Information Signatures

### 8.1 Real proteomes depart from composition-matched i.i.d. baselines

Figure 1 compares real proteomes to matched IID and Markov-1 controls in the (*H*_1_, *I*_1_) plane using bootstrap replicates. By construction, the IID control preserves *H*_1_ but destroys positional dependence, yielding *I*_1_ near zero. Across the 20 proteomes analyzed, the mean adjacent mutual information increased from

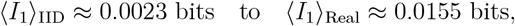

indicating robust short-range dependence beyond composition alone. The mean composition entropy across real proteomes was ⟨*H*_1_⟩_Real_ ≈ 4.164 bits/site (as expected, closely matched by IID controls built from the same *p*_1_).

### 8.2 Information profiles remain elevated beyond Markov-1 decay

To probe dependence across scales, we computed information profiles *I*_*d*_ for separations *d* = 1, …, 25. While Markov-1 controls reproduce *d* = 1 dependence by construction, real proteomes frequently display elevated *I*_*d*_ for *d >* 1 relative to both IID and Markov-1 surrogates. Summarizing dependence beyond adjacency by averaging over *d* = 2, …, 25, we observed:

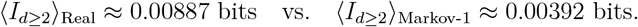

These results indicate dependencies that are not fully explained by a first-order Markov model.

Figure 2 shows representative *I*_*d*_ profiles for three proteomes (human, *E. coli* K-12, and *Drosophila*). Full per-proteome profiles for all 20 datasets can be provided as supplementary figure S1.

**Figure 2:**
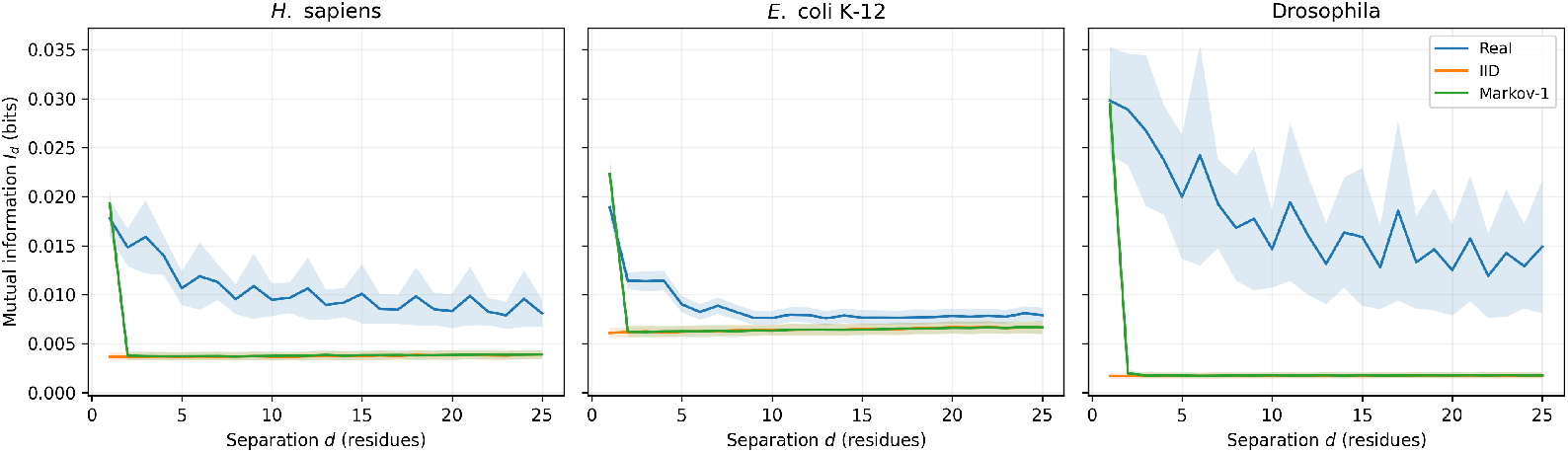
Information profiles *I*_*d*_ (mean ± s.d. across bootstrap replicates) for three representative proteomes. IID controls collapse toward a near-zero baseline for all *d*. Markov-1 controls reproduce the *d* = 1 dependence by construction and decay rapidly thereafter. Real proteomes often exhibit elevated *I*_*d*_ over multiple separations, indicating dependencies beyond a first-order Markov source.

### 8.3 Category-level fingerprints

To summarize cross-clade patterns, Table 1 reports category-averaged signatures: adjacent dependence (*I*_1_), non-adjacent dependence averaged over *d* = 2..25 (*I*_d≥2_), and a compressibility gap relative to IID controls. Categories exhibit distinct statistical fingerprints, consistent with heterogeneous constraints across the tree of life. Importantly, these statistics are descriptive and can reflect a combination of factors including heterogeneous domain composition, repeats, mixture structure, and global constraints rather than any single mechanistic cause.

**Table 1:**
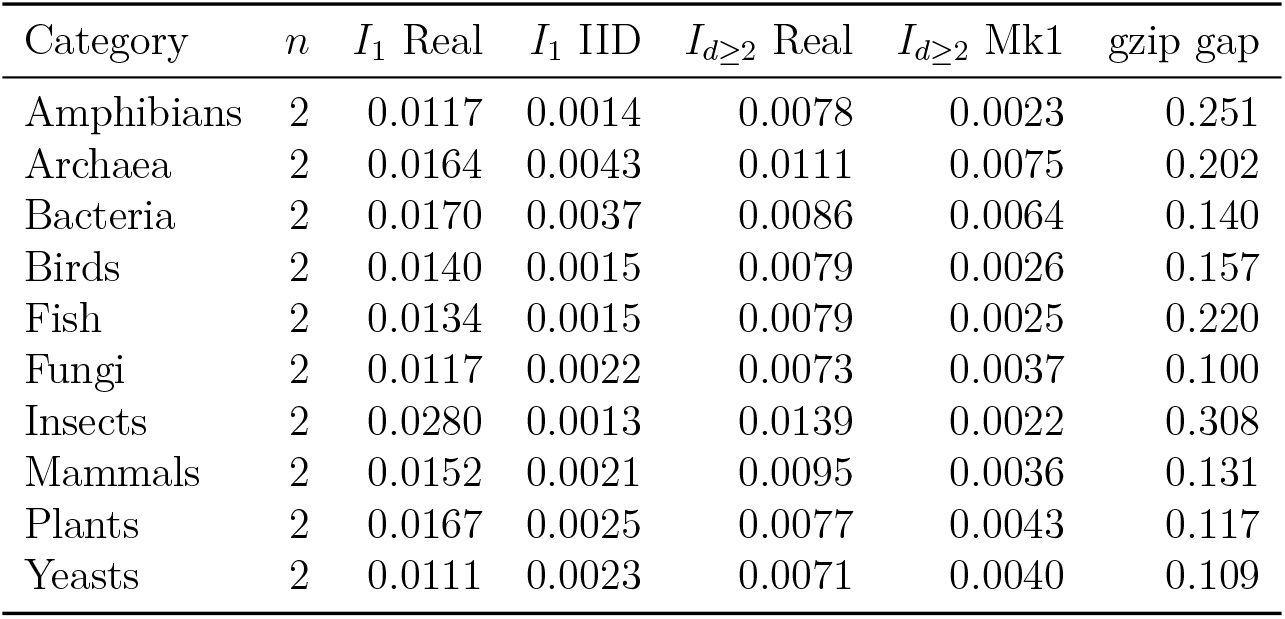
Category-level information signatures (means over two proteomes per category). *I*_*d*≥2_ denotes the mean of *I*_*d*_ over *d* = 2..25. “gzip gap” is Δ*C*_gzip_ = *C*_IID_ − *C*_Real_ (bits/char; positive implies real sequences are more compressible than IID controls).

### 8.4 Compressibility provides an orthogonal redundancy signature

We computed an overhead-corrected gzip bits/character estimate on long concatenations of sampled proteins. At matched composition, IID sequences are expected to be least compressible; deviations indicate redundant patterns and ordering constraints. Across the 20 proteomes, the mean compressibility gap was

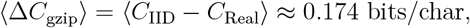

with pronounced gaps in several eukaryotic proteomes. Figure 3 summarizes bootstrap-level gaps.

**Figure 3:**
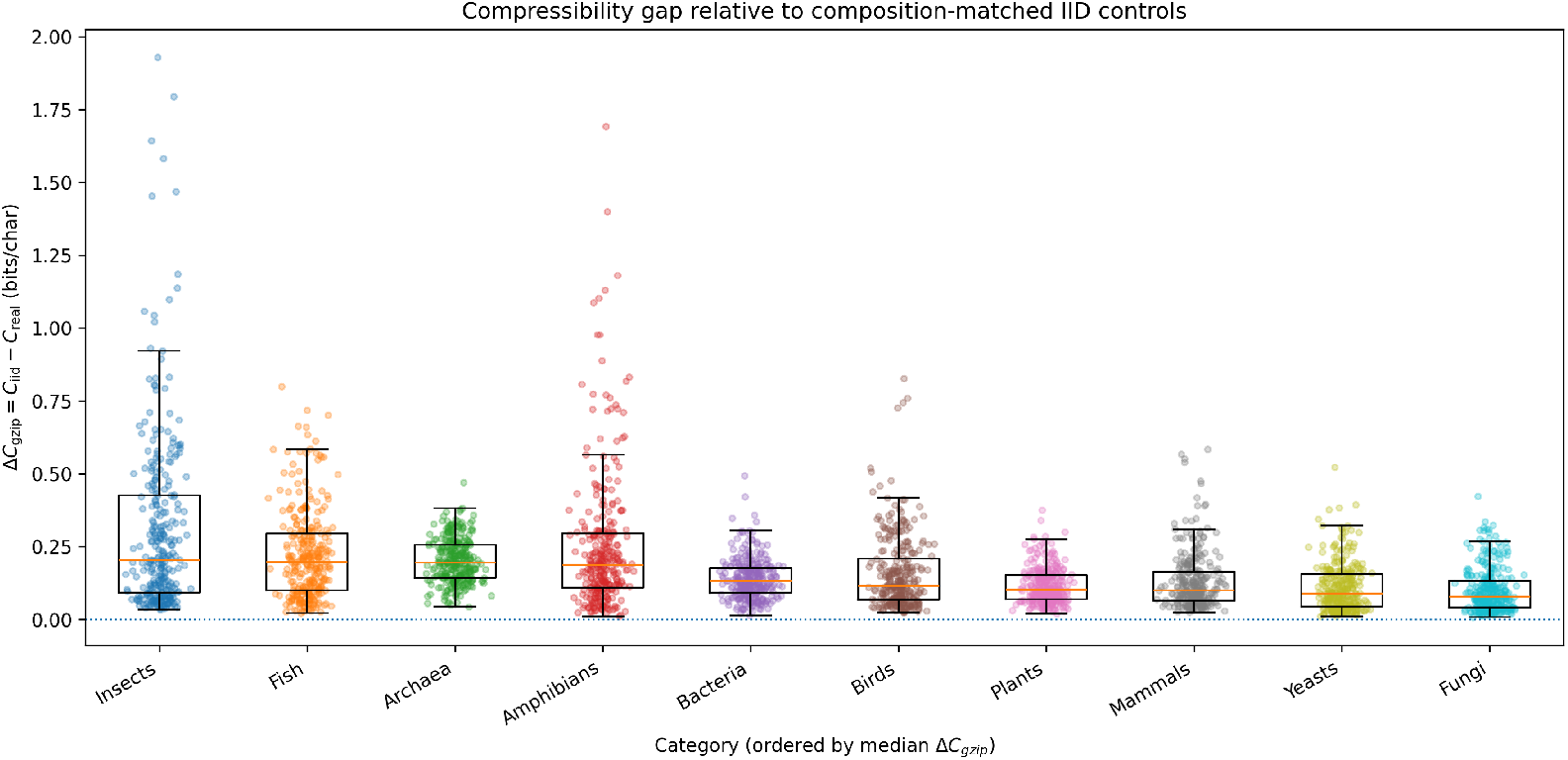
Compressibility gap relative to composition-matched i.i.d. controls, Δ*C*_gzip_ = *C*_iid_ − *C*_real_. Boxplots summarize bootstrap replicates aggregated by broad biological category; points show individual bootstrap replicates. Positive values indicate that real proteome sequences are systematically more compressible than i.i.d. surrogates at matched composition, consistent with non-random ordering constraints.

### 8.5 Entropy-rate reduction from local correlations

Adjacent dependence reduces per-residue uncertainty relative to a composition-only baseline. Using 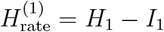, Fig. 4 compares the first-order Markov entropy rate to *H*_1_ across bootstrap replicates. The systematic gap between *H*_1_ and 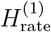 provides an operational measure of local ordering induced by adjacent correlations.

**Figure 4:**
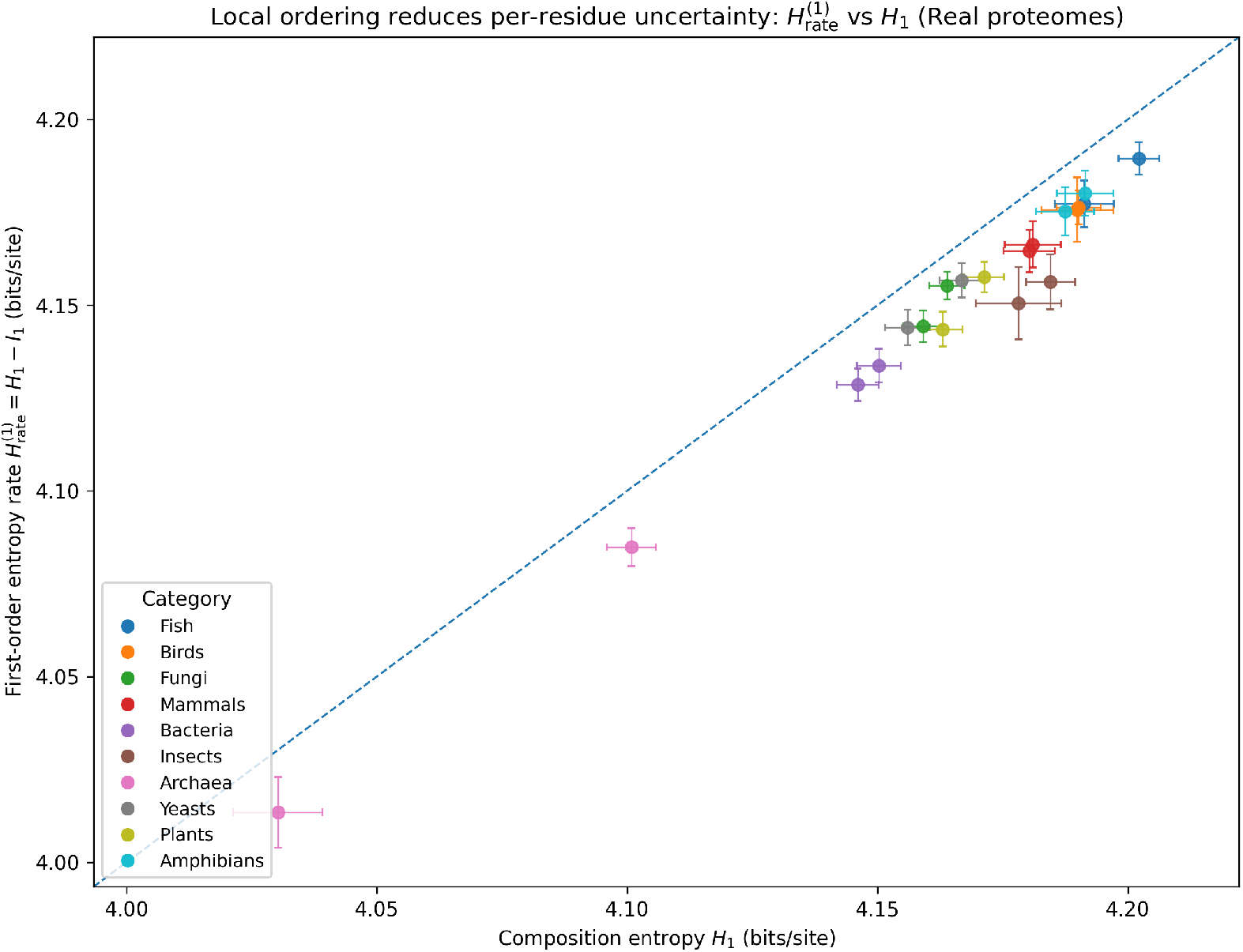
First-order (Markov-1) entropy rate 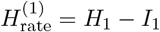 versus composition entropy *H*_1_ for real proteomes (points: proteome-level mean ± s.d. across bootstrap replicates). The diagonal 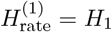 indicates vanishing adjacent dependence (*I*_1_ = 0); systematic displacement below the diagonal quantifies the reduction in per-residue uncertainty induced by local correlations. Dotted lines mark the uniform-alphabet maximum log_2_ 20.

## 9 Discussion and Scope

This work provides a rigorous, model-agnostic language for describing proteomes as constrained stochastic sources over a finite alphabet. The empirical results support three concrete claims.

1. **Proteomes are not i.i.d. at matched composition**. Across 20 proteomes, *I*_1_ increased from ∼ 0.0023 bits (IID controls) to ∼ 0.0155 bits (real), indicating robust local dependence beyond amino-acid composition.
2. **Many proteomes show dependencies beyond first-order Markov structure**. Information profiles *I*_*d*_ frequently remained elevated for *d >* 1 relative to Markov-1 surrogates. This is consistent with constraints not fully captured by adjacent transition statistics. However, such patterns can arise from multiple sources including mixture structure (heterogeneous protein classes), repeats, and domain architecture; therefore, *I*_*d*_ should be interpreted as a descriptive dependence signature rather than direct evidence of a particular mechanistic grammar.
3. **Compressibility provides an orthogonal redundancy signal**. Real proteomes were more compressible than IID controls at matched composition, with mean gap ∼ 0.174 bits/char. This supports the presence of redundant patterns not explained by one-site composition alone.

### 9.1 Relation to maximum-entropy and covariation models

The null-model ladder used here is compatible with a maximum-entropy view: one-site constraints produce IID models, pairwise constraints induce short-range dependencies, and richer constraint sets generically induce longer-range mutual information [3, 6, 7, 18]. The present analysis does not attempt to infer explicit couplings (as in Potts/DCA); instead, it provides a lightweight diagnostic layer for comparing ensembles before committing to mechanistic models.

### 9.2 Limitations

#### Estimation bias

Plug-in estimators for entropy and mutual information can be biased, especially when counts are sparse [13]. Bootstrap variability partially addresses sampling uncertainty but does not eliminate estimator bias. Future work can incorporate bias-corrected or Bayesian estimators.

#### Proteome subsampling and filtering

For computational feasibility, large proteomes were subsampled and sequences shorter than *L*_min_ = 80 were excluded. These choices can shift estimated statistics by changing the mixture of proteins contributing to pooled counts. Sensitivity analyses (varying caps and *L*_min_) are recommended for high-stakes comparisons.

#### Interpretation of long-range dependence

Elevated *I*_*d*_ for *d >* 1 does not uniquely identify a biological mechanism. It can reflect repeats, domain architecture, compositional segmentation, or heterogeneous mixtures of protein families. Disentangling these contributions requires stratified analyses (e.g. by subcellular localization, domain composition, or functional classes).

## 10 Conclusion

We developed a theoretical and empirical framework for viewing proteomes as statistical languages: ensembles of strings generated by constrained stochastic processes. We defined intrinsic descriptors (composition entropy *H*_1_, adjacent mutual information *I*_1_, information profiles *I*_*d*_, and a compressibility proxy), proposed a principled null-model ladder, and proved a general entropy-deficit bound linking rarity constraints to unavoidable statistical structure. Empirically, across 20 UniProt reference proteomes, real sequences depart from composition-matched i.i.d. controls and frequently exhibit dependencies beyond first-order Markov surrogates, yielding distinct cross-clade statistical fingerprints. This framework provides a clean, reproducible foundation for comparing proteome-level sequence structure without requiring structural annotation or semantic assumptions.

