## Supplementary Figure S1 for "Proteins as Statistical Languages: Information-Theoretic Signatures of Proteomes Across the Tree of Life"

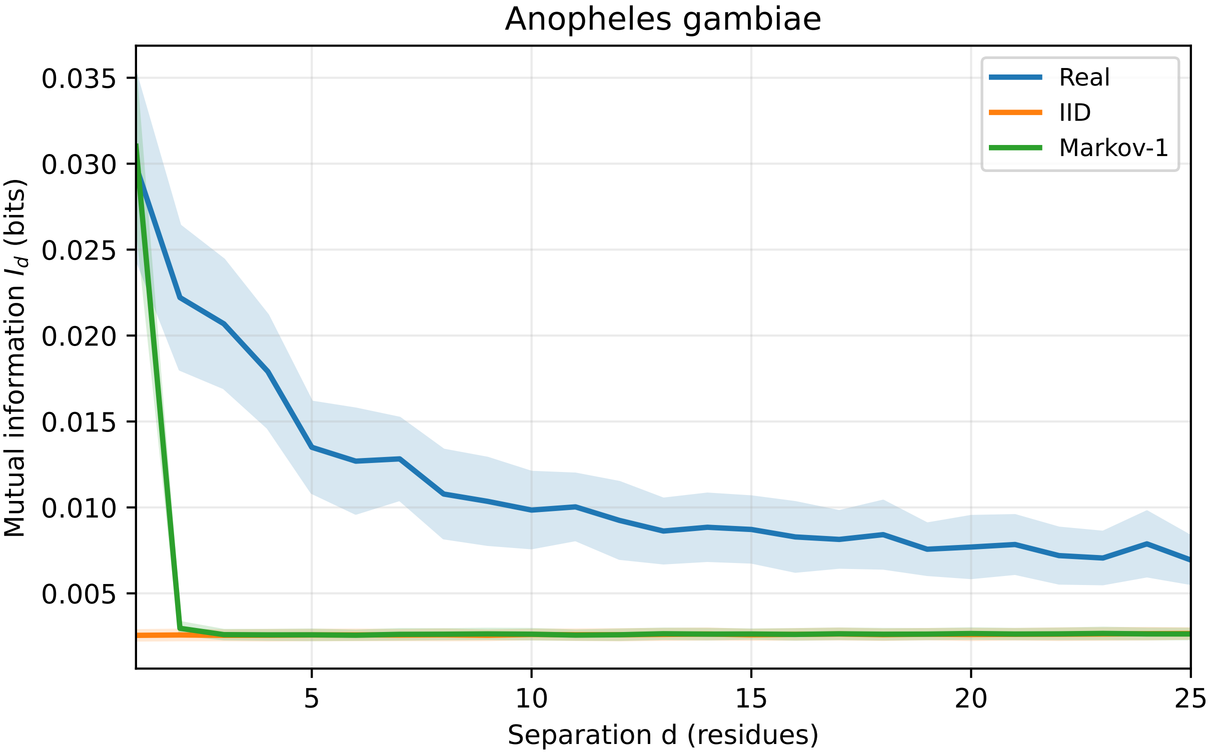

Anopheles gambiae
Real
IID
Markov-1
Mutual information $I_d$ (bits)
0.035
0.030
0.025
0.020
0.015
0.010
0.005
5
10
15
20
25
Separation d (residues)

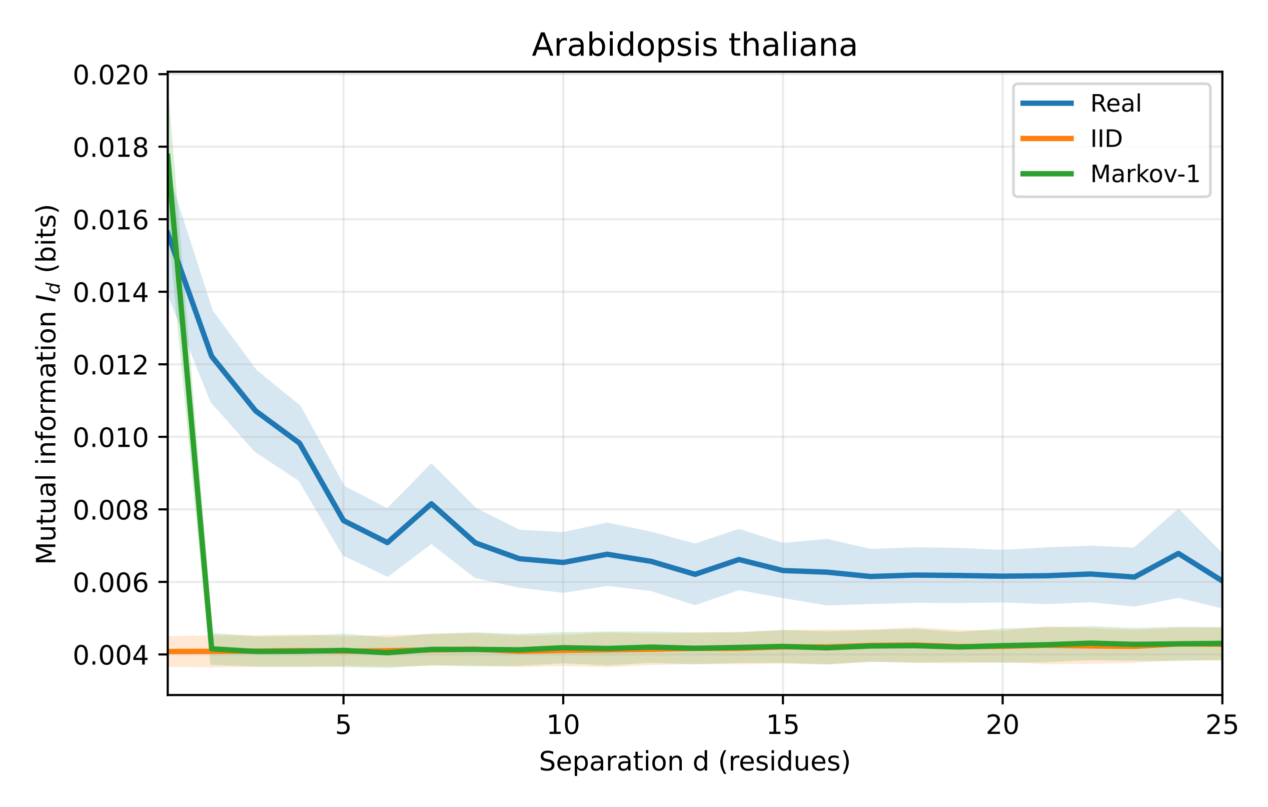
Arabidopsis thaliana
Mutual information $I_d$ (bits)
Separation d (residues)
0.020
0.018
0.016
0.014
0.012
0.010
0.008
0.006
0.004
5
10
15
20
25
Real
IID
Markov-1

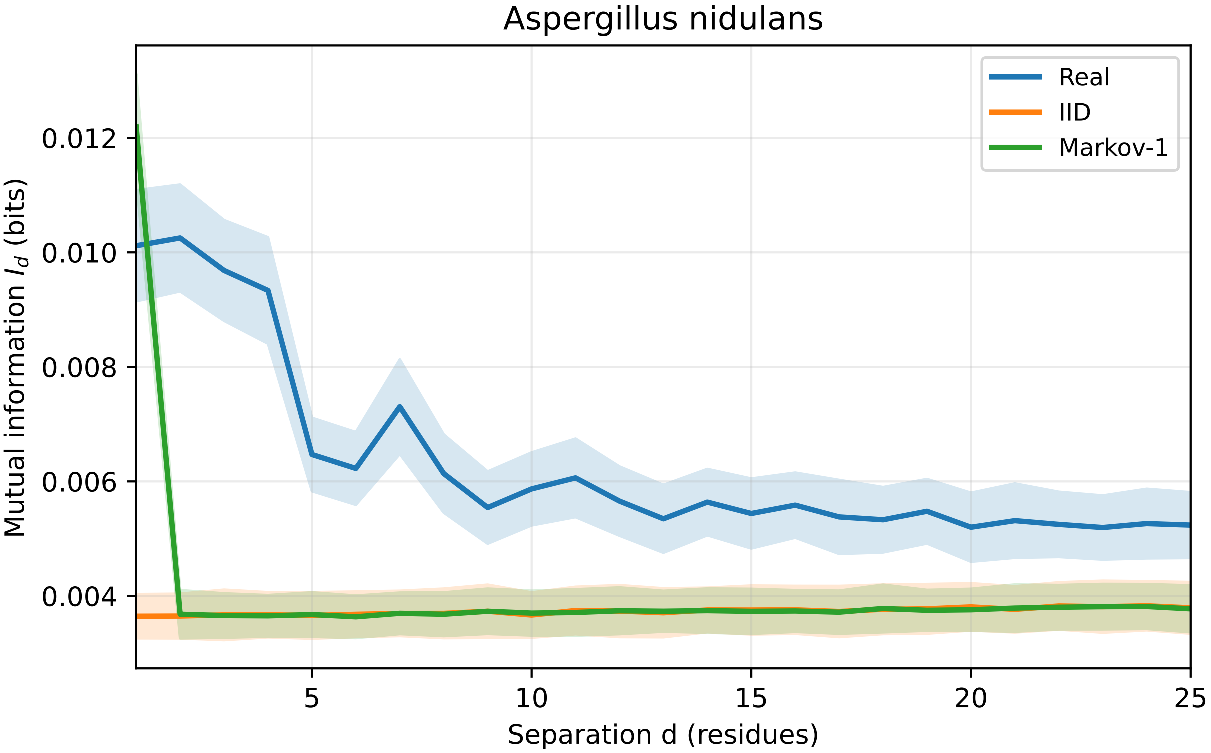

Aspergillus nidulans
Real
IID
Markov-1
Mutual information $I_d$ (bits)
0.012
0.010
0.008
0.006
0.004
5
10
15
20
25
Separation d (residues)

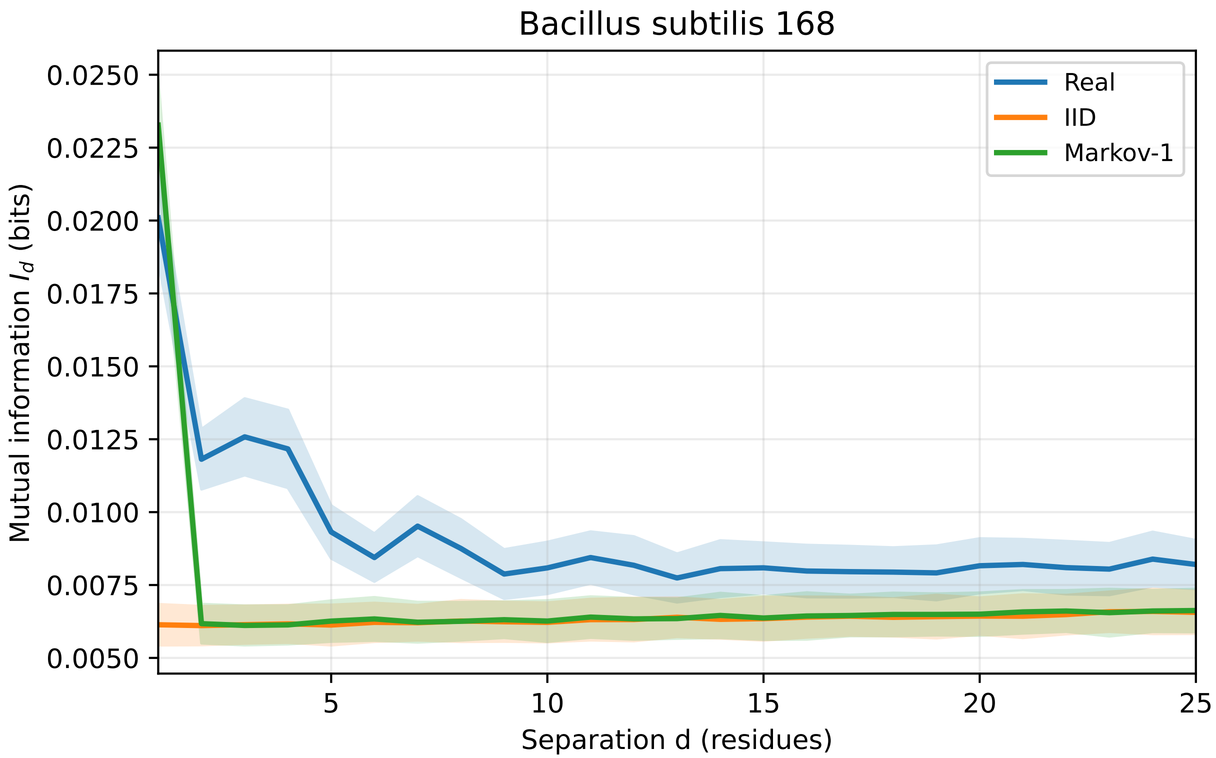

Bacillus subtilis 168
Real
IID
Markov-1
Mutual information $I_d$ (bits)
0.0250
0.0225
0.0200
0.0175
0.0150
0.0125
0.0100
0.0075
0.0050
5
10
15
20
25
Separation d (residues)

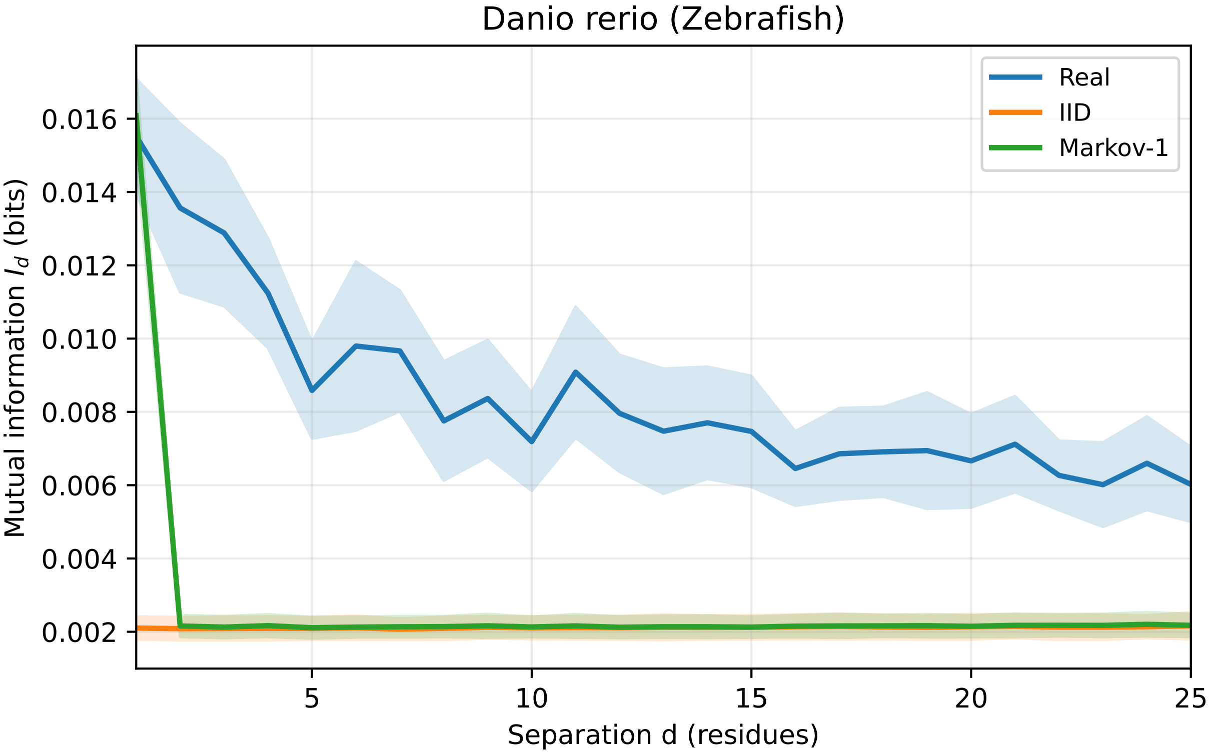
Danio rerio (Zebrafish)
Real
IID
Markov-1
Mutual information $I_d$ (bits)
0.016
0.014
0.012
0.010
0.008
0.006
0.004
0.002
5
10
15
20
25
Separation d (residues)

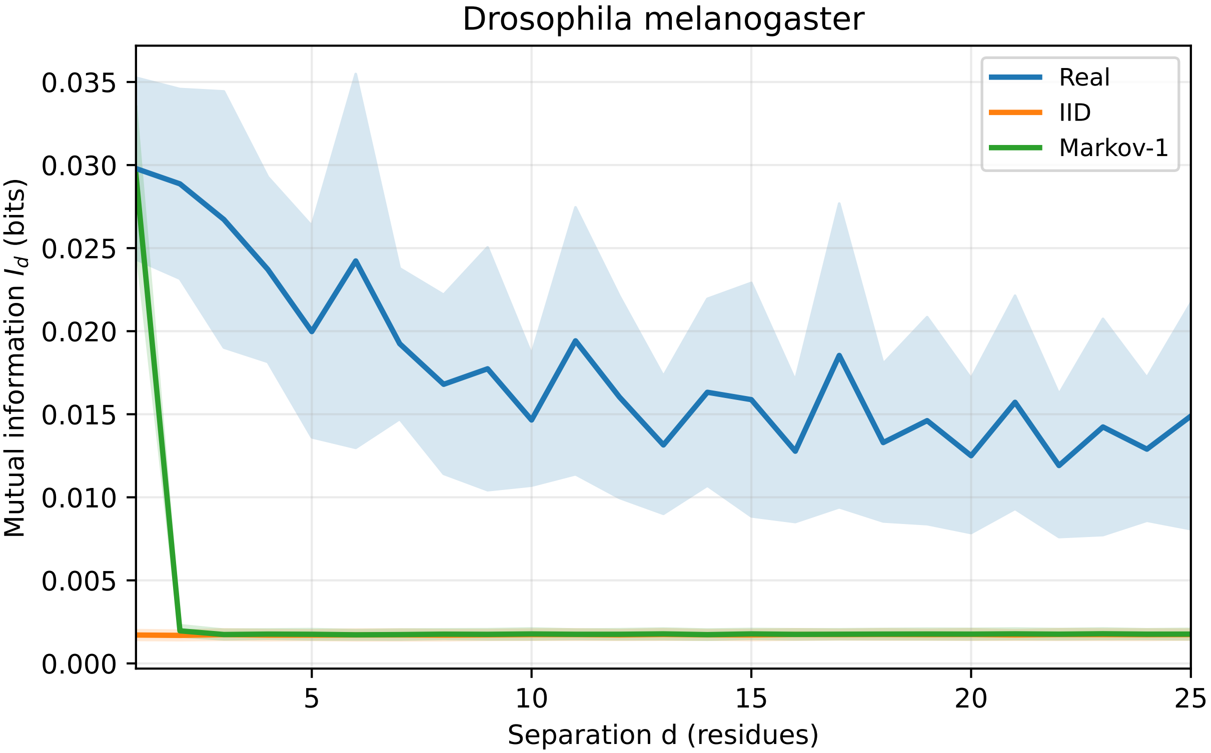

Drosophila melanogaster
Real
IID
Markov-1
Mutual information $I_d$ (bits)
0.000
0.005
0.010
0.015
0.020
0.025
0.030
0.035
5
10
15
20
25
Separation d (residues)

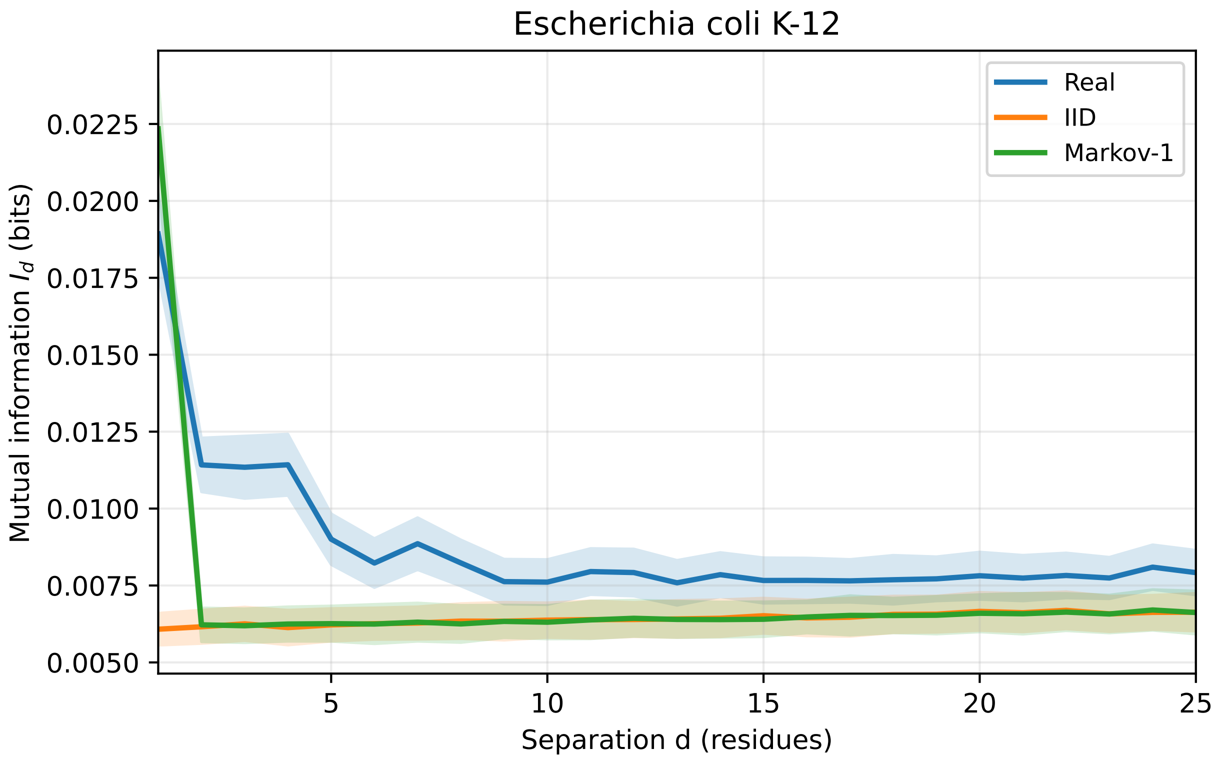
Escherichia coli K-12
Real
IID
Markov-1
Mutual information $I_d$ (bits)
0.0225
0.0200
0.0175
0.0150
0.0125
0.0100
0.0075
0.0050
5
10
15
20
25
Separation d (residues)

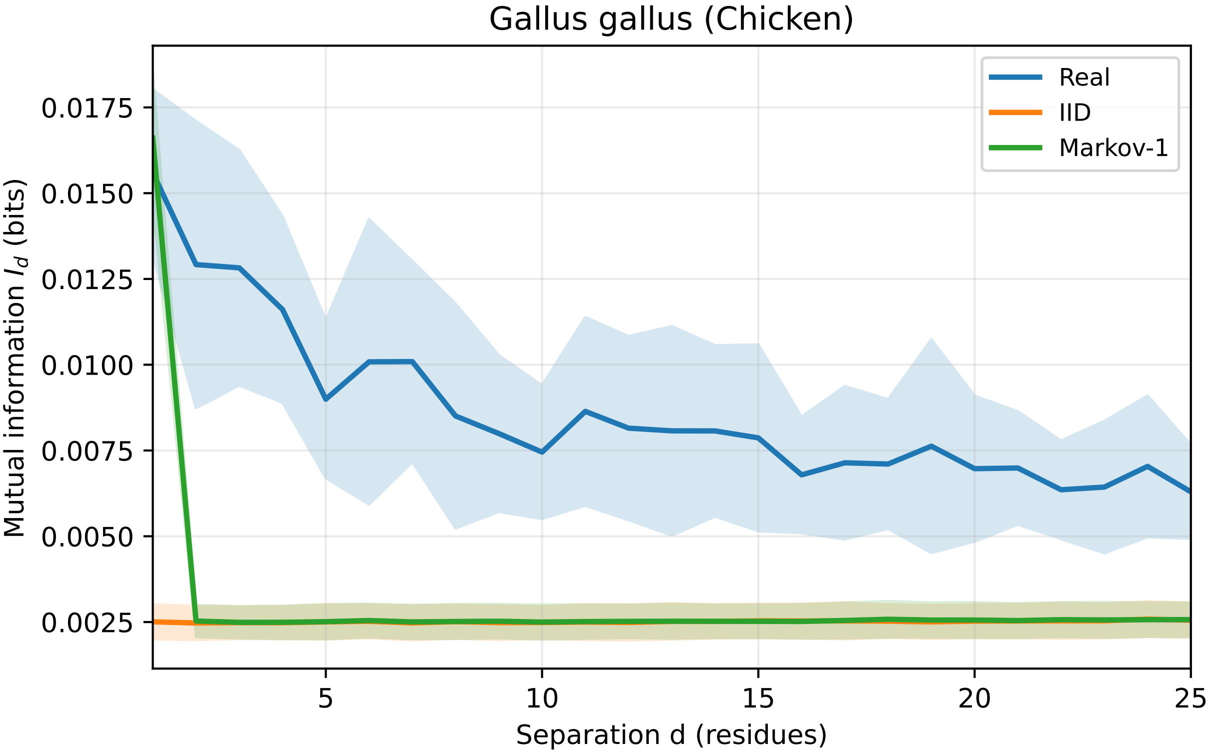
Gallus gallus (Chicken)
Real
IID
Markov-1
Mutual information $I_d$ (bits)
0.0175
0.0150
0.0125
0.0100
0.0075
0.0050
0.0025
5
10
15
20
25
Separation d (residues)

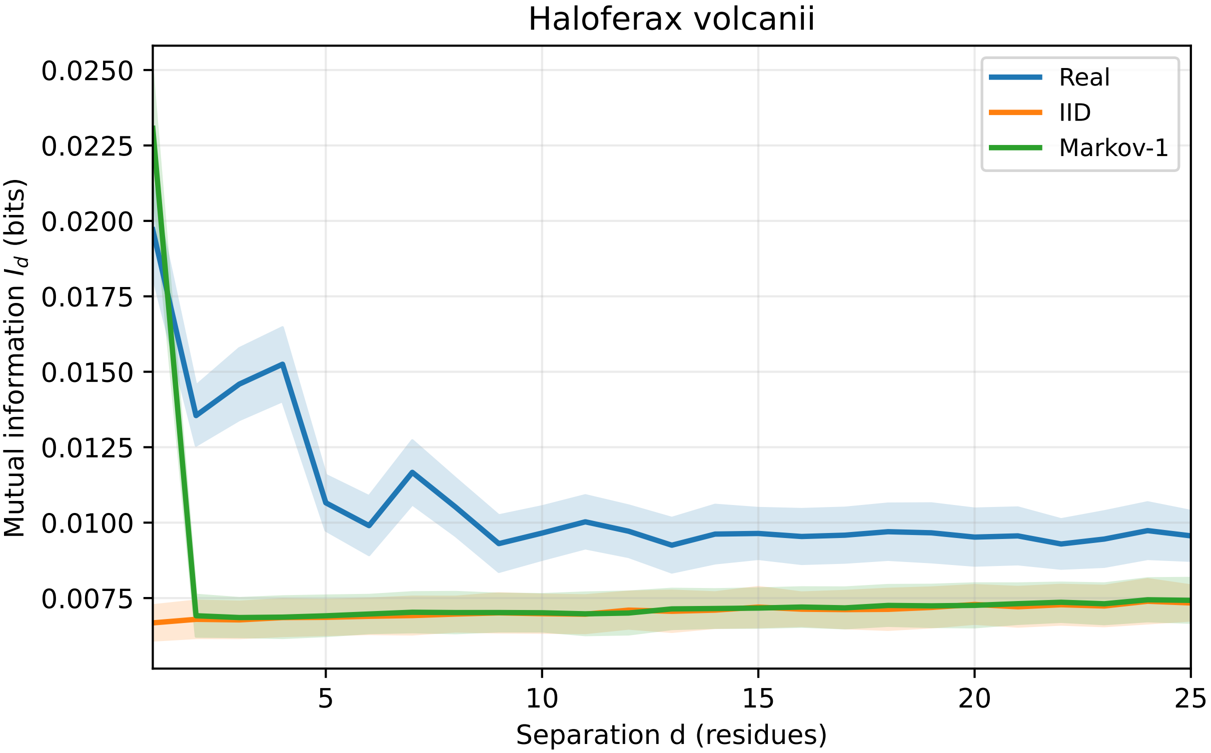
Haloferax volcanii
Real
IID
Markov-1
0.0250
0.0225
0.0200
0.0175
0.0150
0.0125
0.0100
0.0075
Mutual information $I_d$ (bits)
5
10
15
20
25
Separation d (residues)

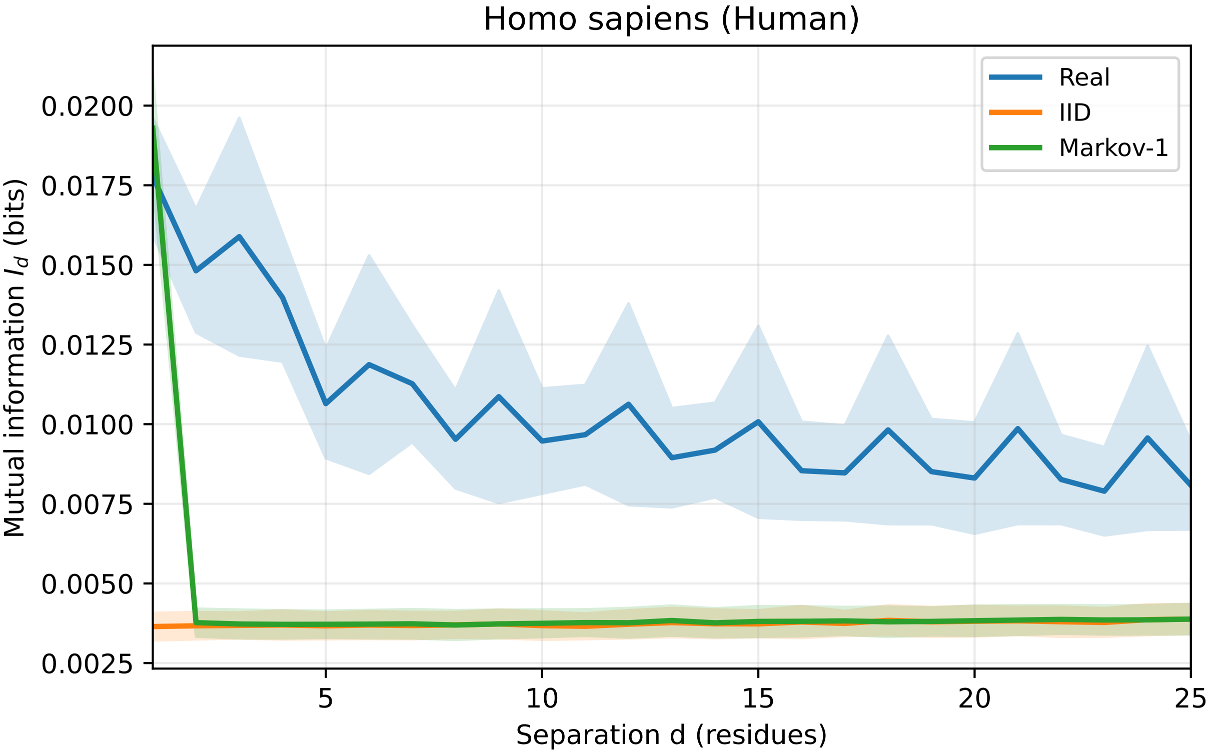
Homo sapiens (Human)
Real
IID
Markov-1
Mutual information $I_d$ (bits)
0.0200
0.0175
0.0150
0.0125
0.0100
0.0075
0.0050
0.0025
5
10
15
20
25
Separation d (residues)

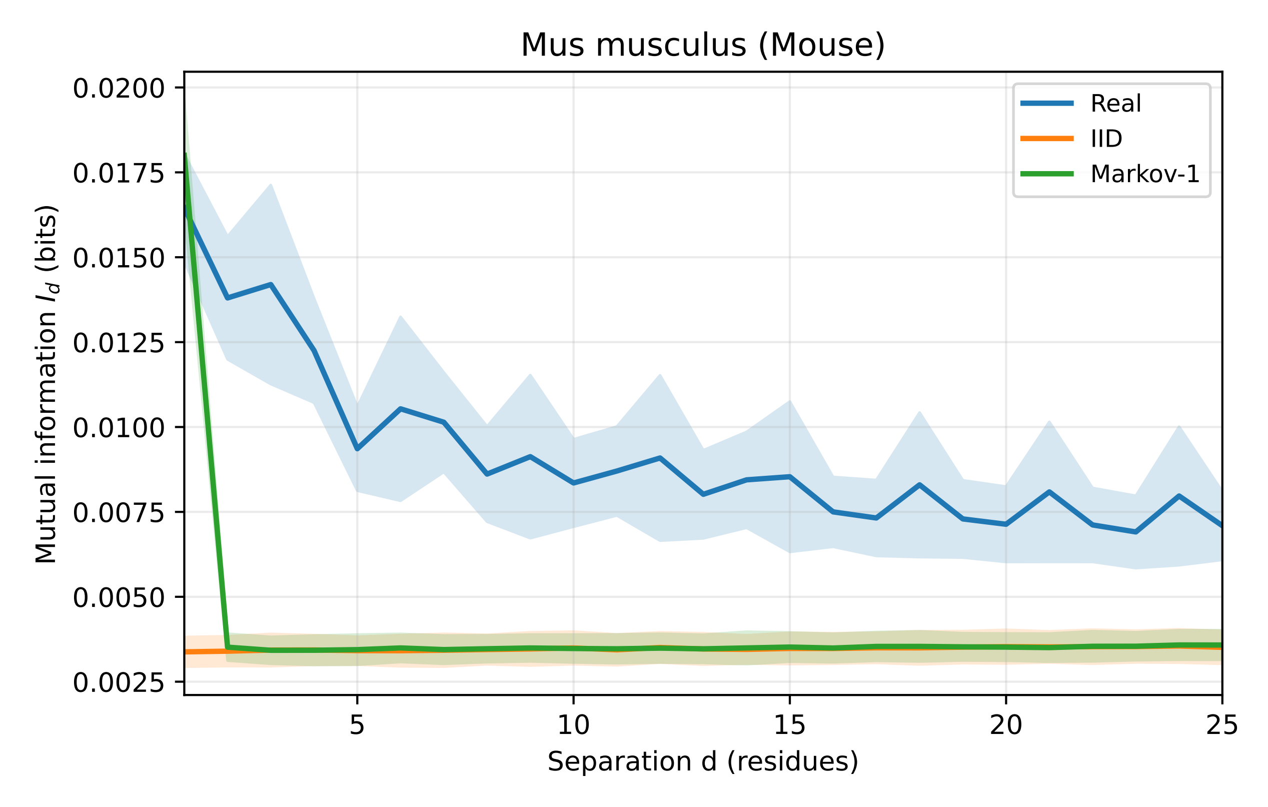

Mus musculus (Mouse)
Real
IID
Markov-1
0.0200
0.0175
0.0150
0.0125
0.0100
0.0075
0.0050
0.0025
Mutual information $I_d$ (bits)
5
10
15
20
25
Separation d (residues)

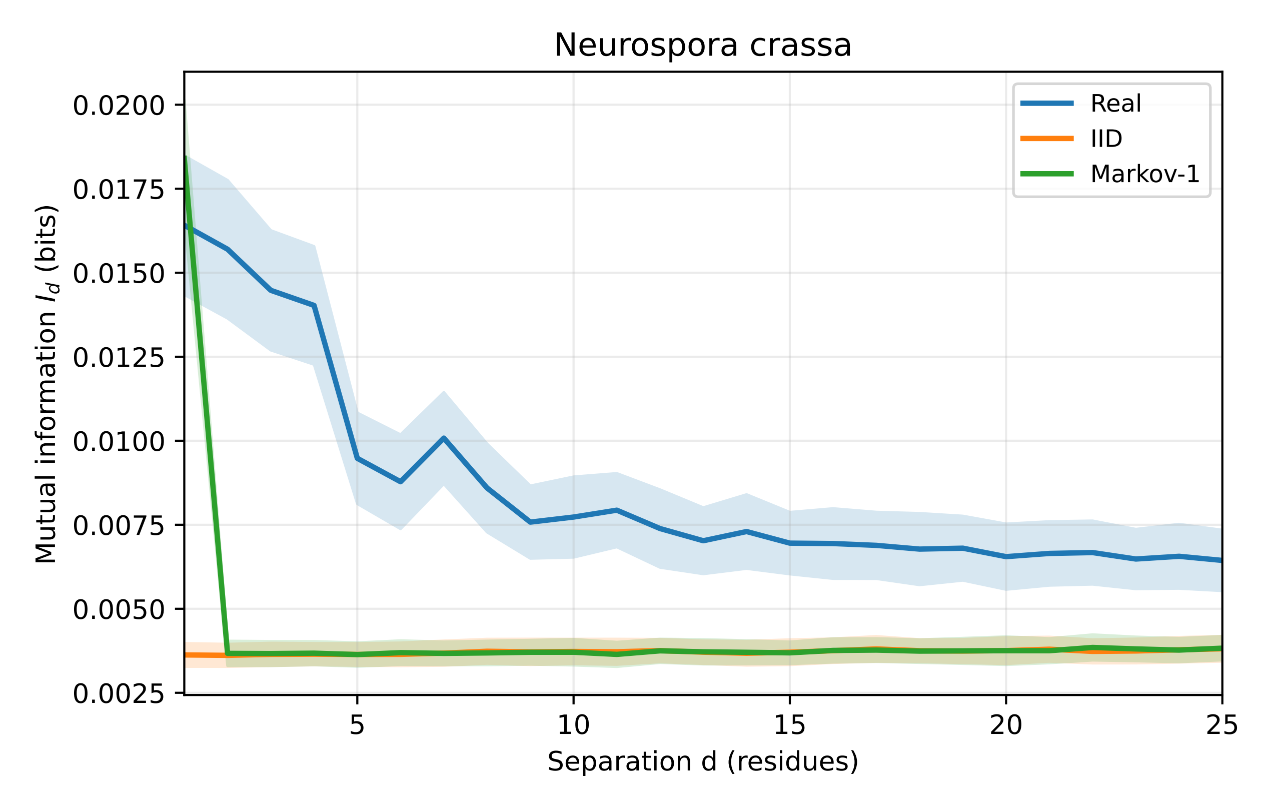

Neurospora crassa
0.0200
0.0175
0.0150
0.0125
0.0100
0.0075
0.0050
0.0025
Mutual information $I_d$ (bits)
5
10
15
20
25
Separation d (residues)
Real
IID
Markov-1

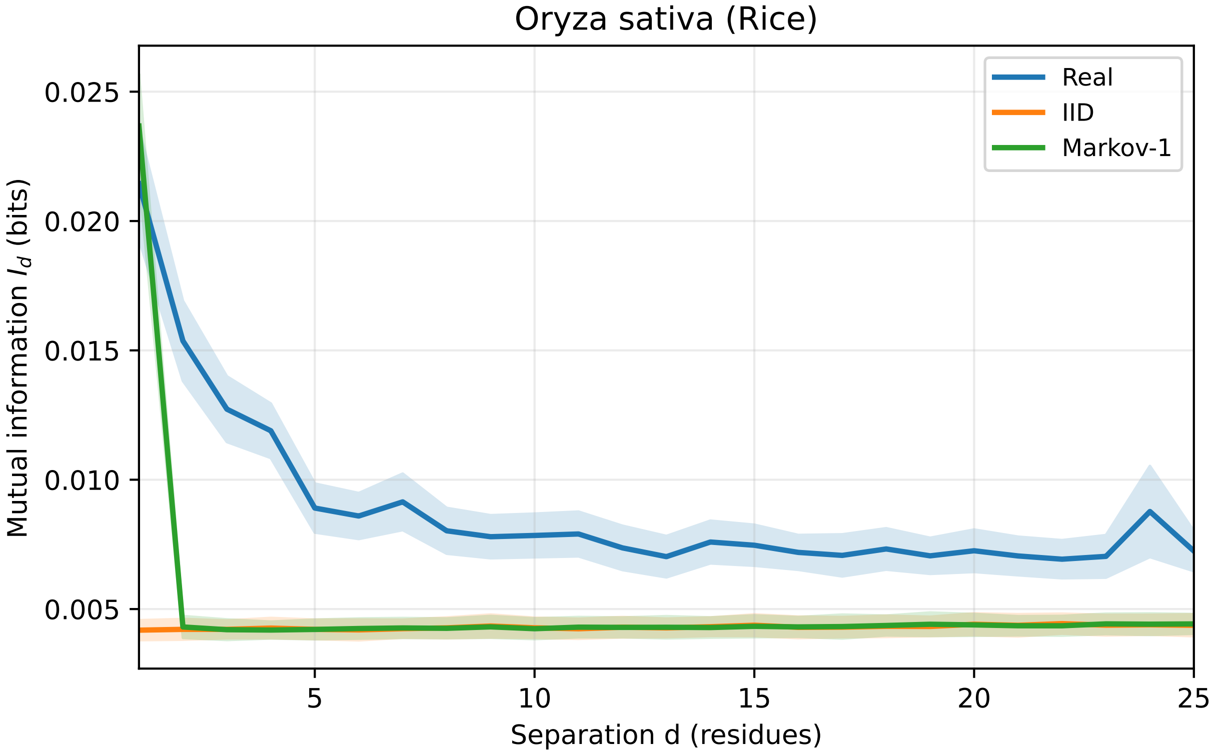

Oryza sativa (Rice)
Real
IID
Markov-1
0.025
0.020
0.015
0.010
0.005
5
10
15
20
25
Mutual information $I_d$ (bits)
Separation d (residues)

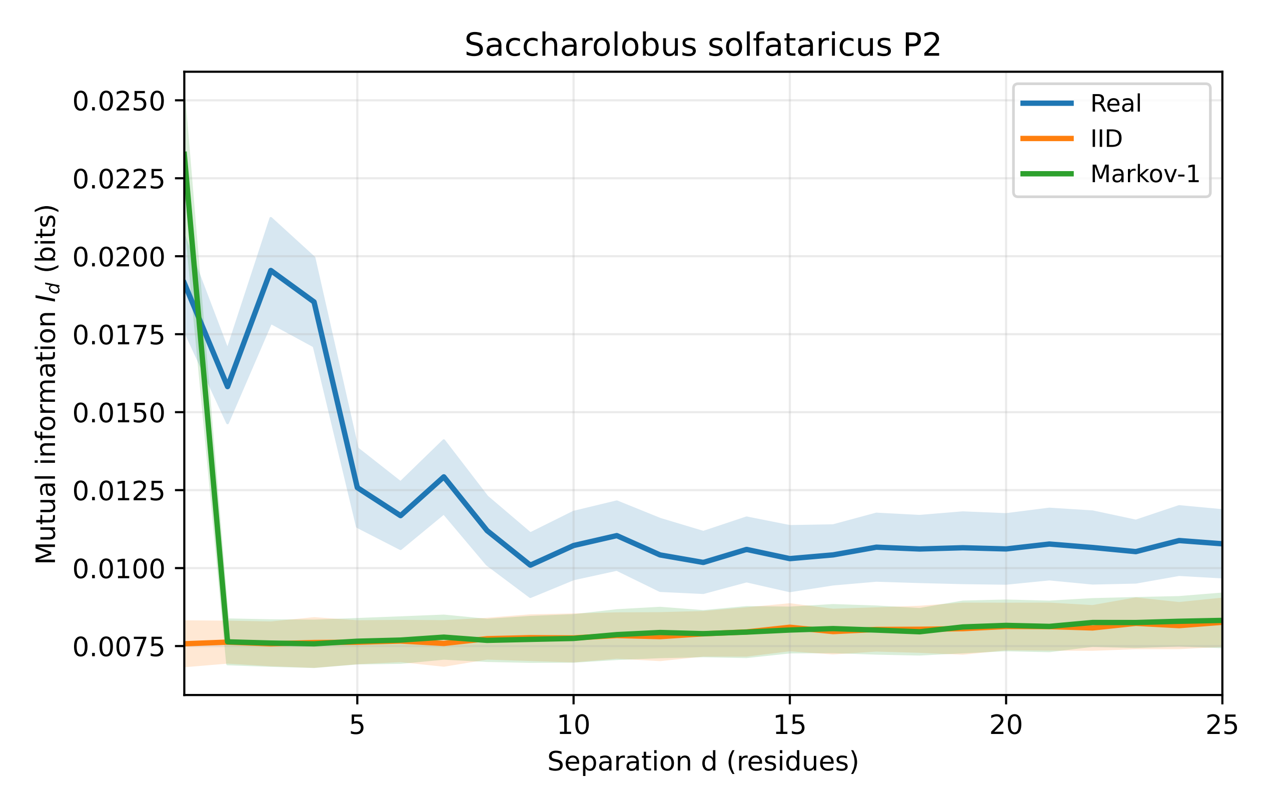
Saccharolobus solfataricus P2
Real
IID
Markov-1
0.0250
0.0225
0.0200
0.0175
0.0150
0.0125
0.0100
0.0075
Mutual information $I_d$ (bits)
5
10
15
20
25
Separation d (residues)

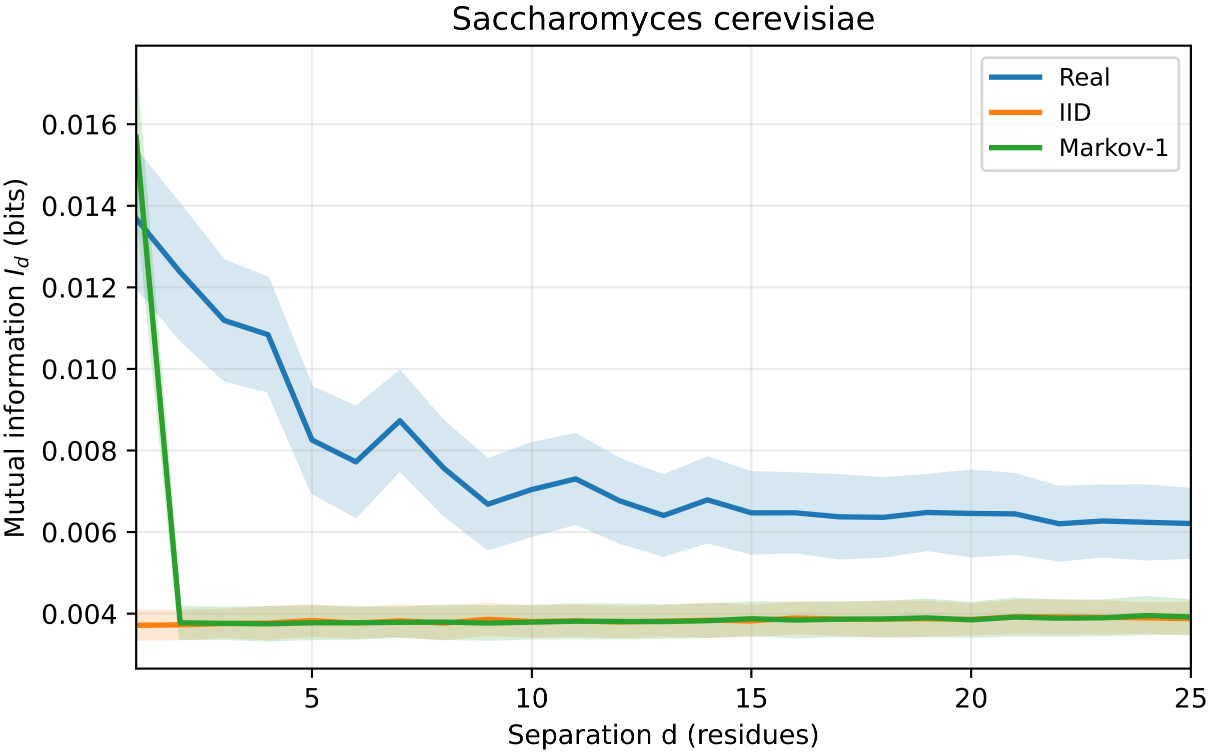

Saccharomyces cerevisiae
Real
IID
Markov-1
Mutual information $I_d$ (bits)
0.016
0.014
0.012
0.010
0.008
0.006
0.004
5
10
15
20
25
Separation d (residues)

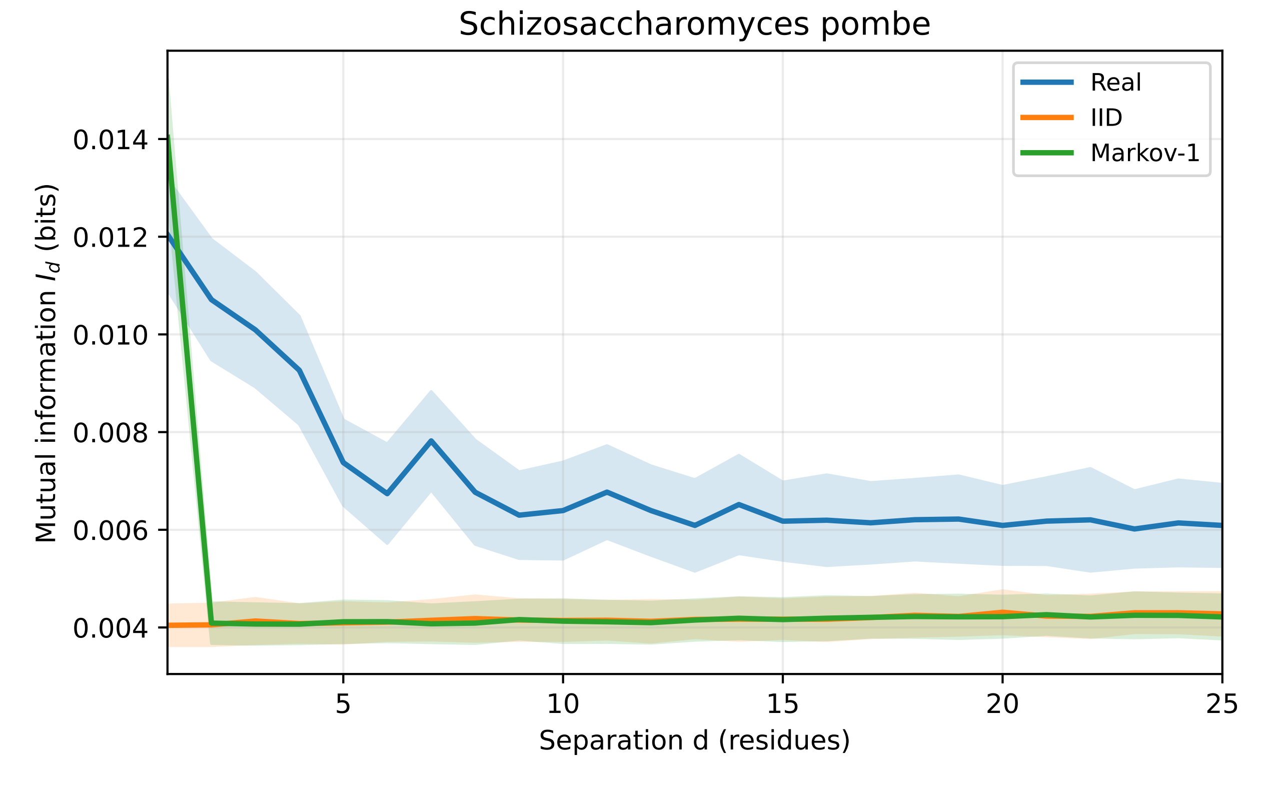
Schizosaccharomyces pombe
Real
IID
Markov-1
0.014
0.012
0.010
0.008
0.006
0.004
Mutual information $I_d$ (bits)
5
10
15
20
25
Separation d (residues)

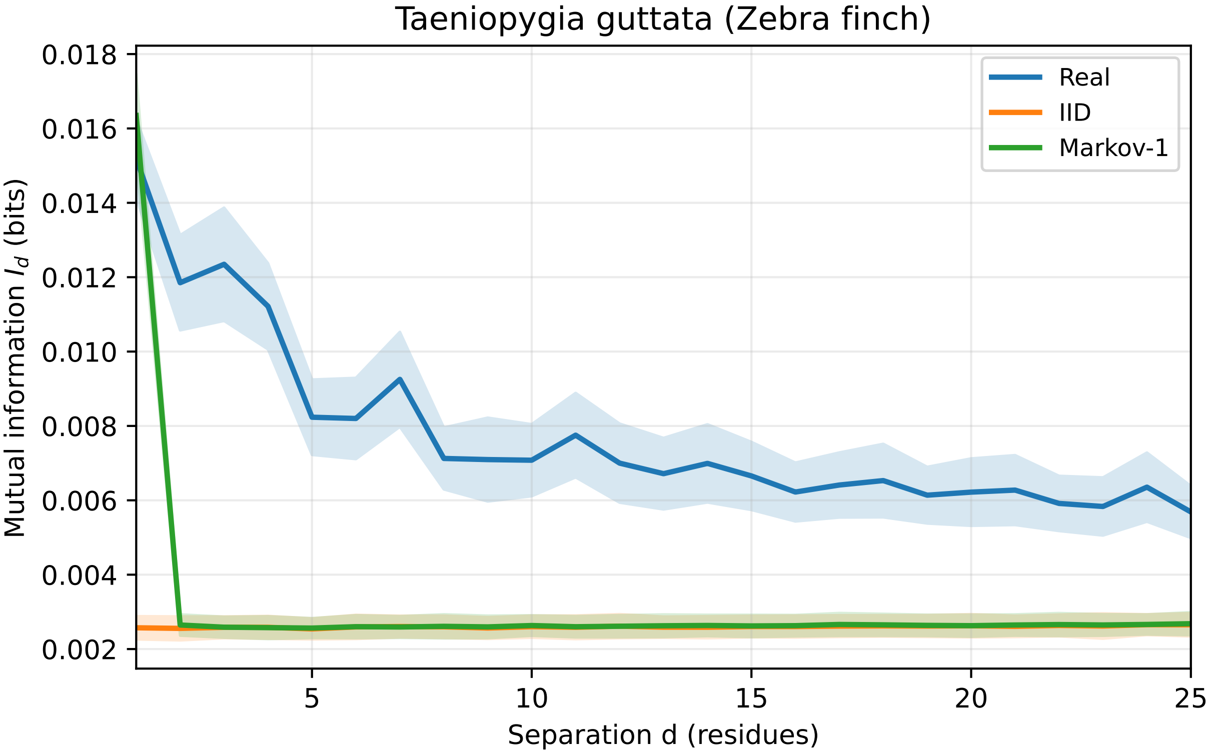

Taeniopygia guttata (Zebra finch)
Real
IID
Markov-1
Mutual information $I_d$ (bits)
Separation d (residues)
0.018
0.016
0.014
0.012
0.010
0.008
0.006
0.004
0.002
5
10
15
20
25

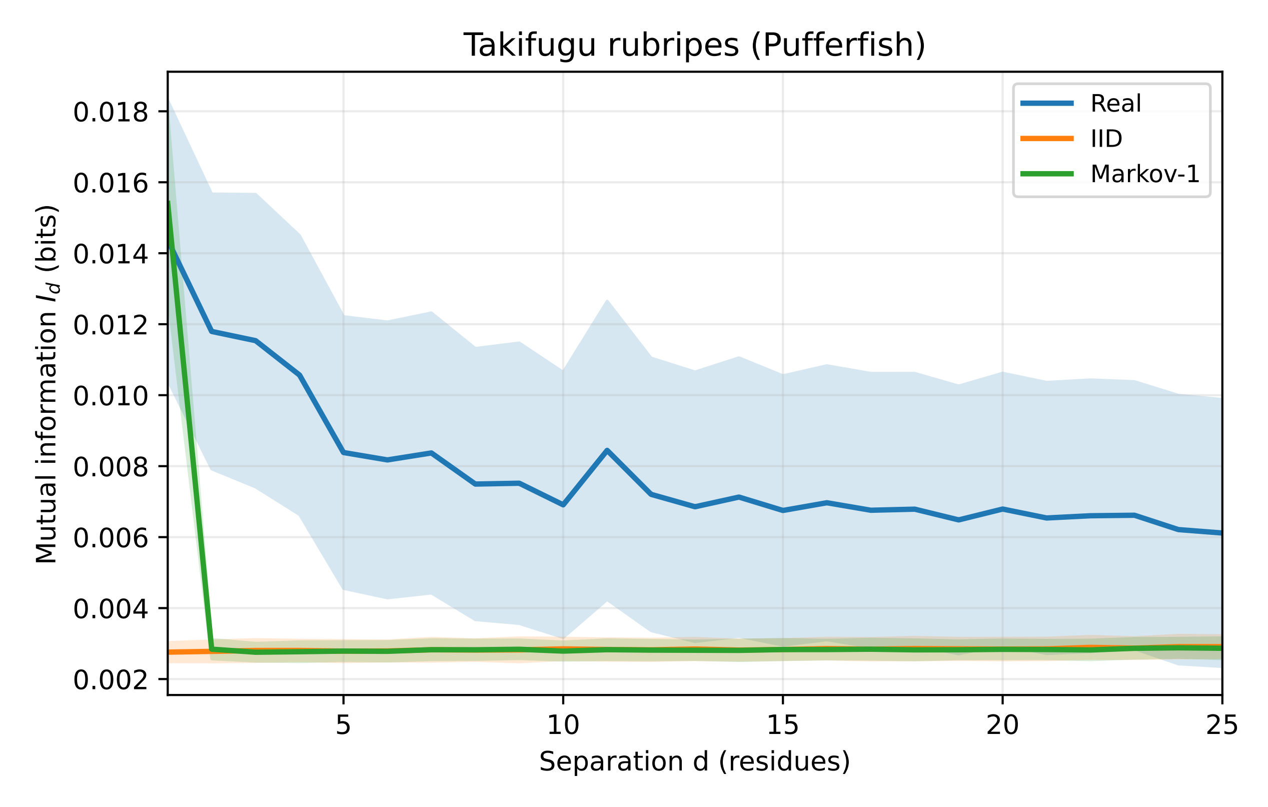

Takifugu rubripes (Pufferfish)
Real
IID
Markov-1
0.018
0.016
0.014
0.012
0.010
0.008
0.006
0.004
0.002
Mutual information $I_d$ (bits)
5
10
15
20
25
Separation d (residues)

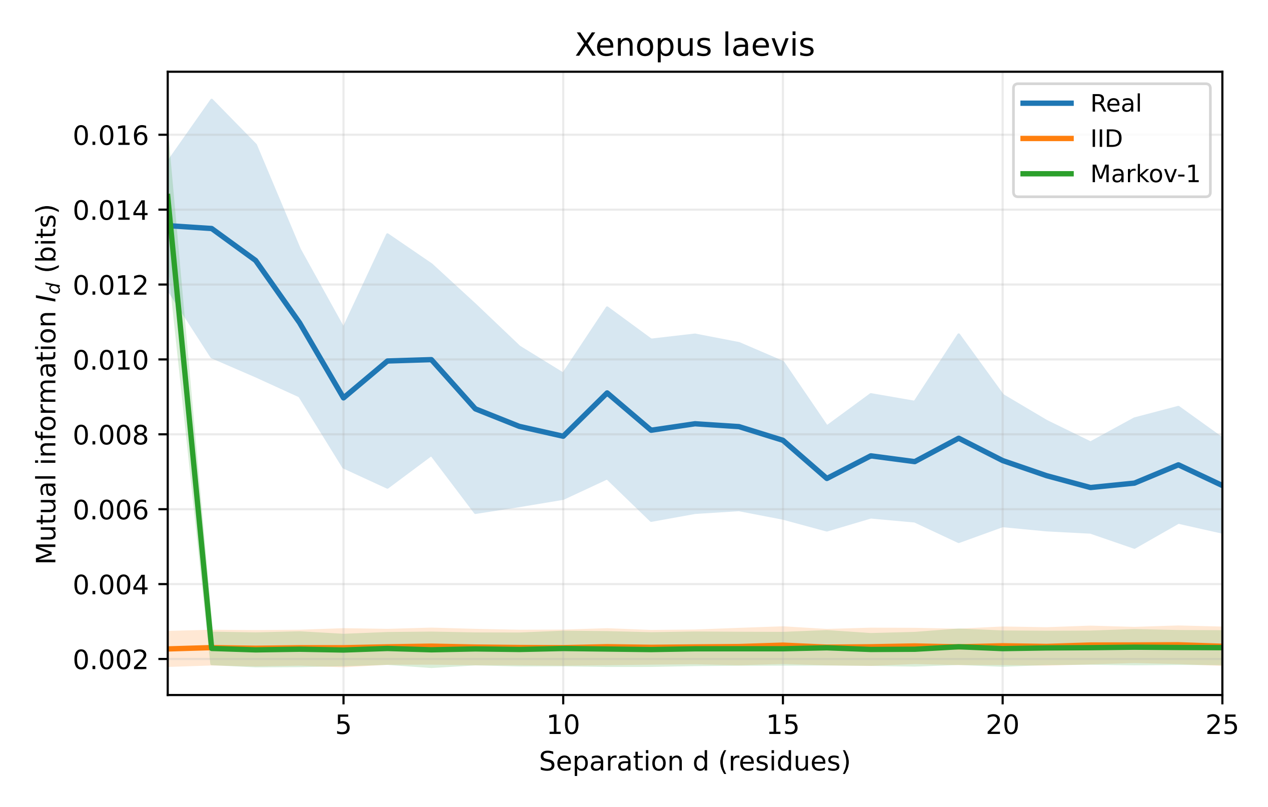

Xenopus laevis
Real
IID
Markov-1
Mutual information $I_d$ (bits)
0.016
0.014
0.012
0.010
0.008
0.006
0.004
0.002
5
10
15
20
25
Separation d (residues)

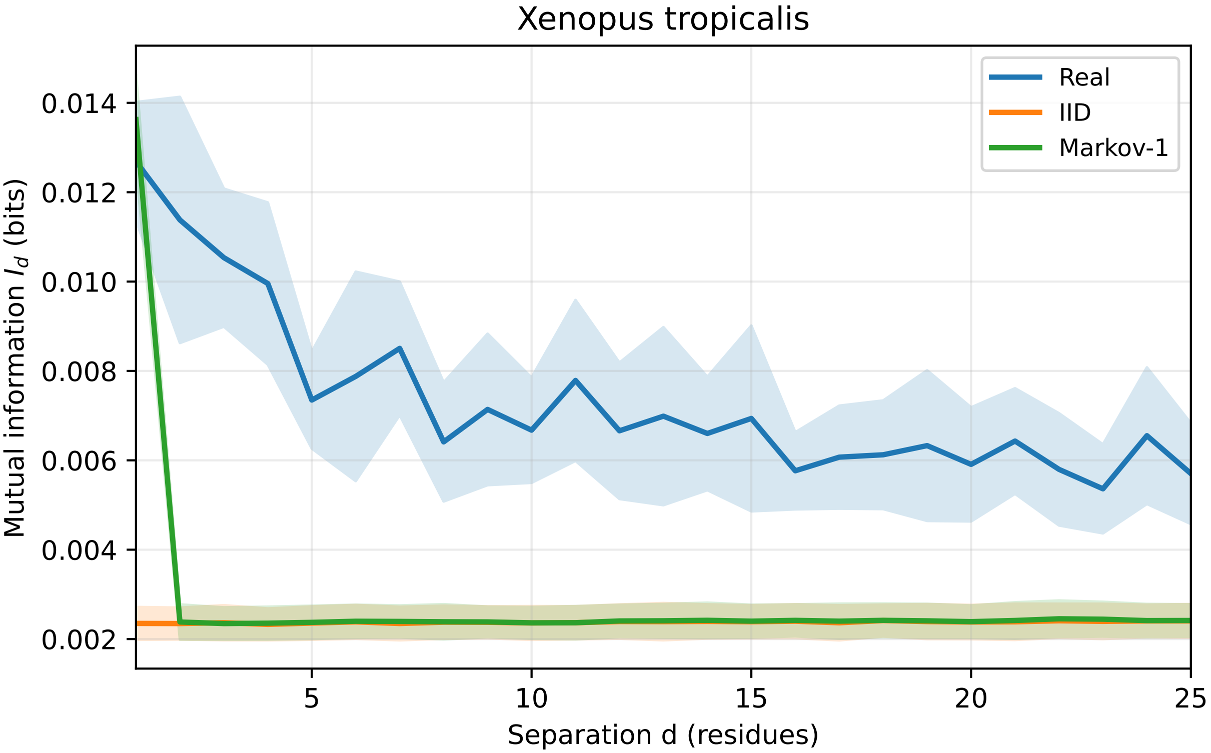
Xenopus tropicalis
Real
IID
Markov-1
0.014
0.012
0.010
0.008
0.006
0.004
0.002
Mutual information $I_d$ (bits)
5
10
15
20
25
Separation d (residues)
